# Cyclosporine A Directly Affects Human and Mouse B cell Migration *in vitro* by Disrupting a HIF-1*α* Dependent, O_2_ Sensing, Molecular Switch

**DOI:** 10.1101/622514

**Authors:** Shannon P Hilchey, Mukta G Palshikar, Dongmei Li, Jessica Garigen, Valantina Cipolla, Juilee Thakar, Martin S Zand

## Abstract

**Background:** Hypoxia is a potent molecular signal for cellular metabolism, mitochondrial function, and migration. Conditions of low oxygen tension trigger regulatory cascades mediated via the highly conserved HIF-1*α* post-translational modification system. In the adaptive immune response, B cells (Bc) are activated and differentiate under hypoxic conditions within lymph node germinal centers, and subsequently migrate to other compartments. During migration, they traverse through changing oxygen levels, ranging from 1-5% in the lymph node to 5-13% in the peripheral blood. Interestingly, the calcineurin inhibitor cyclosporine A is known to stimulate prolyl hydroxylase activity, resulting in HIF-1*α* destabilization and may alter Bc responses directly. Over 60% of patients taking calcineurin immunosuppressant medications have hypo-gammaglobulinemia and poor vaccine responses, putting them at high risk of infection with significantly increased morbidity and mortality.

**Results:** We demonstrate that O_2_ tension is a previously unrecognized Bc regulatory switch, altering CXCR4 chemokine receptor signaling in activated Bc through HIF-1*α* expression and controlling critical aspects of Bc migration. Our data demonstrate that calcineurin inhibition hinders this O_2_ regulatory switch in primary human Bc.

**Conclusion:** This previously unrecognized effect of calcineurin inhibition directly on human Bc has significant and direct clinical implications.

## Background

Calcineurin inhibitors (CNI), such as cyclosporine A (CyA), are used to reduce the risk of allograft rejection in solid organ transplant recipients, and prevent graft-versus-host disease in bone marrow transplantation (1). One side effect of CNI therapy is reduced antigen-specific B cell responsiveness, including hypgammaglobulinemia and reduced protective IgG responses to influenza and other vaccinations (2, 3). This effect has previously been thought to be primarily due to T cell inhibition (4). CNI inhibit calcineurin signaling directly in T cells by preventing calcineurin dependent dephosphorylation of the transcription factor NFAT, blocking nuclear translocation, inhibiting IL-2 production and T cell proliferation (4). In transplant recipients, CNI are thus thought to primarily suppress CD4 T cell help, indirectly causing B cell hypo-responsiveness and depressing antibody production (4). In contrast, direct effects of CNI on B cell function, in particular cellular migration, are few.

Several pieces of evidence suggest that hypoxia may also modulate post-germinal center (GC) B cell migration, and that CNI could interfere with this process. Low tissue oxygen (O_2_) tension (≤1% O_2_; hypoxia) has emerged as an important immune modulating signal (5–10). Intriguingly, low O_2_ tension occurs in secondary lymphoid organs, particularly in draining lymph nodes (LN) and bone marrow (BM) (11, 12). Lymphocytes within these organs express hypoxia induced (transcription) factor-1*α* (HIF-1*α*) at low O_2_ tensions (1% - 4%) (13–15). It has recently been shown that GC hypoxia itself is important for efficient B cell class switching and antibody production in mouse models (16, 17), directly linking hypoxia to B cell function. Despite these results, the mechanism by which B cell function is directly affected by hypoxia during migration to regions of secondary lymphoid tissues[25] with low oxygen O_2_ tension, remains undefined (11, 12).

Although current data support the premise that O_2_ tension modulates transcriptome and molecular signaling events in human T (13, 14, 18–20) and lymphoma cells (21–23), the effects of O_2_ tension has on B cell responses are largely unknown. We have previously demonstrated that HIF-1*α* transcripts are upregulated in both human differentiating B cells *in vitro* and plasma cells migrating *in vivo* through peripheral blood to bone marrow post-vaccination (24, 25).

Coordinated migration of B cells between GC, peripheral blood (PB), spleen and BM is critical for the B cell response (26–29), and is modulated in part by CXCR4 (30) and its ligand CXCL12 (26–29), which are known to be regulated by HIF-1*α* in other cells (13–15). CXCR4 signaling is regulated by transcriptional control, protein expression, and receptor internalization (31). Interestingly, GC B cells have been shown to express surface CXCR4, however, they are unresponsive to CXCL12 signaling (32, 33). As GC B cells encounter O_2_ levels, at times <1%, it is likely that CXCR4 responsiveness is in part controlled by an O_2_ dependent post-translational mechanism, independent of CXCR4 transcription, translation or surface expression. Thus, changes in O_2_ tension as B cells migrate within the GC may directly control the localization and functional activation and differentiation of B cells.

This hypothesis is strongly supported by the O_2_ dependent regulation of several CXCR4 signaling components, including RGS1, which mediates HIF-1*α* induced CXCR4 uncoupling, along with p44/p42 MAPK and MKP-1 (33). Focal adhesion kinase (FAK) is also critical for CXCR4 dependent migration of B cells (15), and is modulated by O_2_ tension in smooth muscle cells (34). In addition, CNI are known to interact directly and indirectly with the HIF-1*α* signaling cascade, and may have a significant role in disrupting the normal hypoxia-induced regulation of B cell migration. For example, CNI destabilize HIF-1*α* in glioma cells by stimulating prolyl hydroxylase activity (35), suggesting CNI have the capacity to disrupt hypoxic responses. Thus, there is strong support for the hypothesis that hypoxia induced pathways are involved in modulation of CXCR4 signaling in B cells and CNI may disrupt these pathways.

Here we demonstrate that migration of human and mouse B cells is regulated by CXCR4 responsiveness via an O_2_ sensing molecular switch, controlled by HIF-1*α*, which is independent of CXCR4 receptor expression or surface localization. Our data support the hypothesis that B cell migration is modulated by CXCR4 chemokine receptor sensitivity, controlled by stabilization of the master regulator HIF-1*α* at low O_2_ levels (<4%), and indeed, we show genetically that HIF-1 *α* is necessary for this effect. Significantly, CyA destabilizes HIF-1*α* in both human and mouse B cells, partially restoring CXCR4 responsiveness at low O_2_ levels. Additional unbiased proteomics data suggests a switch in several metabolic processes potentially facilitating migration. This is consistent with the regulation of extracellular matrix and and intrinsic apoptosis observed in the proteomic analysis. Transient restabilization of HIF-1*α* in CyA treated B cells temporarily restores the O_2_ dependent molecular switch modulating B cell migration. These novel findings have significant implications in regards to CNI suppressed antibody responses, identifying a direct, and potentially therapeutically targetable effect on B cell function, independent of indirect helper T cell effects.

## Methods

### Human subjects protection

This study was approved by the Research Subjects Review Board at the University of Rochester Medical Center (RSRB 71460). Informed consent was obtained from all participants. Research data were coded such that subjects could not be identified, directly or through linked identifiers, in compliance with the Department of Health and Human Services Regulations for the Protection of Human Subjects (45 CFR 46.101(b)(4)).

### Human cell lines and primary B cell isolation

Human cell line: RAMOS (ATCC, Manassas, VA; CRL-1596) Burkitt’s lymphoma B cell line was maintained in complete RPMI 1640 media supplemented with 10% FBS (cR10; Invitrogen, Carlsbad, CA). Primary human B cells: Human peripheral blood was collected by routine phlebotomy in heparinized vacutainers (BD Biosciences, San Diego, CA) and diluted 1:2 with PBS containing 2% FBS. Di-luted blood was then layered over Ficoll-Paque PLUS (GE Healthcare, Chicago, IL) for density centrifugation using Sepmate tubes (Stemcell Technologies, Vancouver, BC, Canada). The lymphocyte layer was then washed twice with PBS (Invitrogen, Carlsbad, CA). PBMCs were then enriched for B cells by using the EasySep Human Pan-B cell Enrichment Kit (Stemcell Technologies, Vancouver, BC, Canada). PBMCs were incubated with an antibody cocktail containing CD2, CD3, CD14, CD16, CD36, Cd42b, CD56, CD66b, CD123, glycophorin A and dextran for 10 minutes. Streptavidin magnetic beads were added to the mix and the sample subjected to magnetic separation. The untouched enriched B cells were transferred to a clean tube, washed twice with PBS. B cells were then analyzed by flow cytometry showing >95% purity of the isolates.

### Mouse splenic B cell isolation

Spleens from euthanized female 9-10 week old C57/Bl6 mice (Taconic Labs, Rensselaer, NY) were removed, mechanically dissociated and cell suspensions passed over a 100-*µ*m nylon cell strainer (BD Biosciences, San Diego, CA). Tissue remaining in the strainer was gently disrupted with the plunger of a 1 mL syringe, and the strainer rinsed with 5 mL of IMDM media (Invitrogen, Carlsbad, CA) containing 200 mM EDTA. The single cell suspension (SCS) was then passed over a 70-*µ*m nylon cell strainer (BD Biosciences, San Diego, CA) to further remove clumps. Excess RBCs were removed by magnetic positive selection (Imag; BD Biosciences, San Diego, CA) after incubation of the cell suspension with 50 *µ*L Ter119 magnetic beads (BD Biosciences, San Diego, CA) per 1 × 10^7^ total cells at 4 °C for 30 min. Unbound cells were transferred to new tubes and washed once with PBS with 2% FBS and 0.4% EDTA. B cells were enriched from the SCS using Mouse Pan-B Cell Isolation kits (Stemcell Technologies, Vancouver, BC, Canada). Briefly, cells were suspended at 1 × 10^8^ cells/mL in PBS with 2% FBS and 0.4% EDTA. Normal Mouse serum was added at 50 *µ*L/mL followed by the Mouse Pan-B cell Isolation Cocktail at 50 *µ*L/mL. Cells were incubated for 10 minutes at RT. RapidSpheres, at 50 *µ*L/mL, were added and incubated for 5 min. at RT. Samples were resuspended to a volume 2.5 mL with PBS with 2% FBS and 0.4% EDTA and placed on an Imag (BD Biosciences, San Diego, CA) for 5 min. Unbound cells were transferred to new tubes; flow cytometry indicated >95% purity of the isolates.

### O_2_ controlled cell treatments

RAMOS B cells or human and mouse primary B cells were resuspended in cR10 media at 5 × 10^4^ cell/mL and added to appropriate sized tissue culture plates or flasks depending on total volume. To select flasks, CyA and/or DMOG (Sigma-Aldrich, St. Louis, MO) was added at the concentration indicated in the text and/or figure legends. Plates or flasks were then placed within controlled environment C-chamber 37 °C, 5% CO_2_ incubator inserts (Biospherix, Parish, NY) that were equilibrated O/N at the indicated O_2_ levels or placed within a traditional 5% CO_2_, 37 °C incubator (19% O_2_). Cells were incubated for 24 hours, removed and subjected to either the chemotaxis, western blot or proteomics assays.

### Chemotaxis assay

The human B cell line, RAMOS cells, or magnetic bead enriched primary human peripheral blood or mouse splenic B cells were resuspended in freshly prepared migration media (PBS + 1% BSA), and 5 × 10^5^ cells (in 75 *µ*L) loaded into the upper chambers of 5 *µ*m polycarbonate 96 well transwell plates (Corning, Corning, NY). In triplicate wells, migration media containing varying concentrations of CXCL12 was added to the bottom chambers of the transwells (225 *µ*L per well). Plates were incubated for 1 hour at appropriate O_2_ levels in the C-chamber incubator inserts (Biospherix, Parish, NY), at 37 °C and 5% CO_2_. Upper transwells were removed and 100 *µ*L of Cell Titer Glo (Promega, Madison, WI) added to each bottom well and incubated at RT for 10 min. Luminescence was readout on a SynergyTM HT microplate reader (BioTek Instruments, Winooski, VT) and relative luminescent units (RLU) reported. Supplementary Figure 3 demonstrates the direct correlation between RLU and cell numbers, through a 4-*log*_10_ range, independent of O_2_ levels.

### Western bloting

Nuclear lysates were prepared from B cell pellets using NE-PER nuclear and cytoplasmic extraction kits and protein concentration determined using micro BCA protein assay kits, both according to the manufacture’s recommendations (Thermo Fisher Scientific, Waltham, MA). Nuclear lysates (10*µ*g per lane) were resolved on precast NuPAGE 4-12% Bis-Tris protein gels (Invitrogen, Carlsbad, CA) and proteins transferred to PVDF membranes (Bio-Rad, Hercules, CA). The membrane was then blocked with 1X TBS-T (Tris-Buffered Saline (Bio-Rad, Hercules, CA)) with 0.05% Tween-20 (Sigma-Aldrich, St. Louis, MO) + 5% blotting grade nonfat dry milk (Bio-Rad, Hercules, CA) for 1 hour at RT. Blots were washed 3 × with TBS-T and then primary antibody diluted as indicated in TBS-T + 5% milk. For human B cell blots a 1:250 dilution mouse anti-human HIF-1*α* (BD Biosciences, San Diego, CA) or for mouse blots a 1:100 dilution goat anti-mouse HIF-1*α* wsa added and the blots incubated O/N at 4 °C. As a loading control, blots were also probed with a 1:200 dilution mouse anti-actin primary antibody and the blots incubated O/N at 4 °C. For detection of the human HIF-1*α* or actin blots, a 1:1,000 dilution, in PBS + 5% milk, of horseradish peroxidase (HRP) conjugated goat anti-mouse Ig secondary was used. For mouse HIF-1*α*, a 1:2,000 dilution HRP conjugated donkey anti-goat IgG was used. Secondaries were added and the blots incubated for 1 hour at RT. Blots were were washed 3 X with TBS-T, rinsed with distilled water and freshly prepared ECL substrate (Thermo Fisher Scientific, Waltham, MA) added and the blots were imaged on a ChemiDoc MP imaging system. Densitometry was performed using Image Lab software version 6.0.1 (Bio-Rad, Hercules, CA).

### Proteomics sample preparation

RAMOS cells were resuspended in cR10 media at 5 × 10^5^ cell/mL at a final volume of 10 mL in T25 tissue culture flasks (BD Biosciences, San Diego, CA) and either left untreated or cyclosporine A (Sigma-Aldrich, St. Louis, MO) added to a final concentration of 1 *µ*g/mL. Pairs of untreated or CyA treated cells were then incubated at either 19% or 1% O_2_ levels for 24 hours. Cells were then rapidly harvested into 50 mL conical tubes (BD Biosciences, San Diego, CA) containing 30 mL of ice cold PBS. Cells were pelleted and washed 3 × 10 mL ice cold PBS. After the final wash, cells were resuspended in 1 mL ice cold PBS and transferred to a 1.5 mL microcentrifuge tubes and pelleted. Supernatants were removed and the cell pellets flash frozen in liquid nitrogen. Cell pellets were stored at −80°C until samples were lysed for analysis.

Lysis buffer consisted of 5% SDS, 50 mM triethylammonium bicarbonate (TEAB). Cell lysis or each pellet was done by adding 100 *µ*L of lysis buffer per 10^6^ cells, followed by vortexing and sonication using a QSonica sonicator. Sonication cycles consisted of 5 × 10 second sonications with one minute incubations on ice between each cycle. Samples were centrifuged for 5 minutes at 16,000 × g, and the supernatant was collected. Protein concentration was determined by BCA (Thermo Fisher Scientific, Waltham, MA). 25 *µ*g of protein from each sample was removed, and brought up to 25 *µ*L in lysis buffer. Disulfide bonds were reduced by addition of dithiothreitol (DTT) to 2 mM, followed by incubation at 55 °C for 1 hour. Alkylation was performed by adding iodoacetamide to 10 mM and incubating at room temperature in the dark for 30 minutes. 12% phosphoric acid was added to a final concentration of 1.2%, followed by the addition of 6 volumes of 90% methanol, 100 mM TEAB. The resulting solution was added to S-Trap micros (Protifi, Huntington NY), and centrifuged at 4,000 × g for 1 minute. The STraps were washed twice with 90% methanol, 100 mM TEAB. 20 *µ*L of 100 mM TEAB containing 1 *µ*g of trypsin was added to the S-Trap, followed by an additional 20 *µ*l of TEAB.

Samples were placed in a humidity chamber at 37 °C and were allowed to digest overnight. The S-Trap was centrifuged at 4,000 × g for 1 minute to collect the digested peptides. Twenty *µ*L of 0.1% trifluoroacetic acid (TFA) in acetonitrile was added to the S-Trap, and one more centrifugation step was done, and the solutions were pooled, frozen, and dried down in a Speed Vac (Labconco, Kansas City, MO). Tandem mass tag (TMT) ten-plex reagents (0.2 mg) (Thermo Fisher Scientific, Waltham, MA) were removed from −20 °C and allowed to reach room temperature prior to dissolving each tag in 20 *µ*L of acetonitrile. Samples were re-constituted in 25 *µ*L of TEAB, the TMT tags were added to the samples, and incubated at room temperature for one hour. The reaction was quenched by the addition of 3 *µ*L of 5% hydroxylamine. 20% (5 *µ*g) of each sample was combined, frozen, and dried in the Speed Vac.

To increase coverage, samples were fractionated using C18 spin columns. Columns were conditioned with acetonitrile, followed by equilibration with 100 mM ammonium formate (AF), pH 10. The samples were resuspended in 50 *µ*L of AF and added to the spin columns. After washing the columns with water and then the AF buffer, samples were eluted with 10%, 12.5%, 15%, 17.5%, 20%, 22.5%, 25%, and then 50% acetonitrile in the AF buffer. These fractions were then frozen, dried down, and resuspended in 0.1% TFA in water, and placed into autosampler vials.

### Protein data acquisition

Peptides were injected onto a 30 cm C18 column packed with 1.8 *µ*m beads (Sepax), with an Easy nLC1000 HPLC (Thermo Fisher Scientific, Waltham, MA), connected to a Q Exactive Plus mass spectrometer (Thermo Fisher Scientific, Waltham, MA). Solvent A was 0.1% formic acid in water, while solvent B was 0.1% formic acid in acetonitrile. Ions were introduced to the mass spectrometer using a Nanospray Flex source operating at 2 kV. The gradient began at 6% B and held for 2 minutes, increased to 30% B over 85 minutes, increased to 50% B over 10 minutes, then ramped up to 70% B in 4 minutes and was held for 5 minutes, before returning to starting conditions in 4 minutes and re-equilibrating for 10 minutes, for a total run time of 120 minutes at a 300 nL/minute flow rate. The Q Exactive Plus was operated in data-dependent mode, with a full scan followed by 10 MS/MS scans. The full scan was performed over a range of 400-1700 m/z, with a resolution of 70,000 at m/z of 200, an automatic gain control (AGC) target of 10^6^, and a maximum injection time of 50 ms. Peptides with a charge state between 2-5 were picked for fragmentation. Precursor ions were fragmented by higher-energy collisional dissociation (HCD) using a collision energy of 35 and an isolation width of 1.0 m/z. MS2 scans were collected with a resolution of 35,000, a maximum injection time of 120 ms, and an AGC setting of 1e^5^. The fixed first mass for the MS2 scans was set to 110 m/z to ensure TMT reporter ions were always collected. Dynamic exclusion was set to 25 seconds.

### Proteomics data analysis

Raw data was searched using the SEQUEST search engine within the Proteome Discoverer software platform, version 2.2 (Thermo Fisher Scientific, Waltham, MA), using the SwissProt human database (36). Trypsin was selected as the enzyme allowing up to 2 missed cleavages, with an MS1 mass tolerance of 10 ppm, and an MS2 mass tolerance of 0.025 Da. Carbamidomethyl on cysteine, and TMT on lysine and peptide N-terminus were set as a fixed modifications, while oxidation of methionine was set as a variable modification. Percolator was used as the false discovery rate (FDR) calculator, filtering out peptides which had a q-value greater than 0.01.

Reporter ions were quantified using the Reporter Ions Quantifier node, with an integration tolerance of 20 ppm, and the integration method being set to “most confident centroid”. Protein abundances were calculated by summing the intensities of the reporter ions from each identified peptide, while excluding any peptides with an isolation interference of 30% or more. Low abundance proteins with less than one count per experiment/replicate were removed resulting in 5048 proteins. Proteins’ abundance was *log*_2_-transformed, and normality was confirmed by diagnostic plots such as quantile-quantile plot. The differential expression analysis was performed using the implementation of the empirical Bayes statistic in the R package *limma* (37) across all conditions. Differentially expressed (DE) proteins are defined as p-value of ≤ 0.01 and a *log*_2_-fold change of ≥ 0.5. Pathway analysis of DE proteins was performed using the hypergeometric test and 1392 canonical gene sets curated in MSigDB (38).

### Proteome network analysis

To evaluate the associations among proteins and neighborhood of DE proteins, we assembled a co-expression network using 3919 proteins with coefficient of variation ≥ 0.01. The edges were weighted by the absolute Pearson correlation coefficient between the connected nodes. Each edge in the coexpression network was assigned a z-score by permutation of sample labels 50 times. We retained only edges with a weight (Pearson correlation coefficient) that had an absolute z-score of ≥ 1.5, i.e., the real correlation was at least 1.5 standard deviations away from the mean of correlations calculated by perturbing sample labels. Edges with a weight ≤ 0.5 were removed. This resulted in a network of 1408 nodes and 1211 edges. The implementation of the edge-betweenness community detection algorithm in the R package *igraph* (39, 40) was used to identify communities with ≥ 10 genes. We performed a gene set enrichment analysis using the hypergeometric test. To further map genes/proteins known to be functionally related to HIF1A and CXCR4, 33 pathways from KEGG (41), Biocarta (http://www.biocarta.com/genes/allpathways.asp) and Reactome (42) were identified. The compiled list of 1504 unique genes from these pathways was used in the STRING database (43) to build a network where the edges are weighted based on a score calculated based on strength of experimental evidence of interaction, homology, co-expression, and database annotations. This network is referred to as the *knowledge based network*.

### Statistical Methods

Permutation tests within the general linear models framework were used to fit the data to compare differences groups. Group comparisons of interest were obtained by pairwise comparisons using Tukey’s method to control for the multiple testing error rate. We adjusted for CXCL12 concentrations in the models when HIF-1*α* levels were compared across multiple CXCL12 concentrations. The statistical analyses were conducted using the *lmPerm* and *multcomp* packages within the statistical analysis software R version 3.5.1 (44, 45). The significance level for all tests was set at 5%.

## Results

### Human and mouse B cell CXCR4 hypo-responsiveness is induced by low O_2_ levels and this correlates with HIF-1*α* stabilization

In order to examine B cell CXCR4 responsiveness at different O_2_ levels, we developed a novel high throughput *in vitro* transmigration assay system that combines a 96 well transwell plate format with a rapid luminescent readout of migratory cell numbers. This assay was coupled with precise O_2_ level control using C-Chamber O_2_ control incubator chambers (Biospherix, Parish, NY) to measure cell migration under varying levels of hypoxia. As is evident from Figure 1A, the RAMOS human B cell line migrates in response to a CXCL12 chemokine gradient at 19% O_2_ in a dose dependent fashion. However, incubation of these B cell lines for 24 hours at 1% or 4% O_2_ results in significantly decreased migration (Figure 1A). Importantly, we observed no effect on RAMOS viability, proliferation or surface CXCR4 expression (Supplementary Figure 1) irrespective of O_2_ levels, suggesting that the decrease in chemotaxis is not due to decreased cellular viability and/or proliferation, nor due to loss of CXCR4 surface expression. In addition, this chemotaxis assay is highly reproducible and the decreased chemotactic activity observed at lower O_2_ levels is highly significant (*p*<0.001 for all pairwise comparisons).

**Fig. 1.**
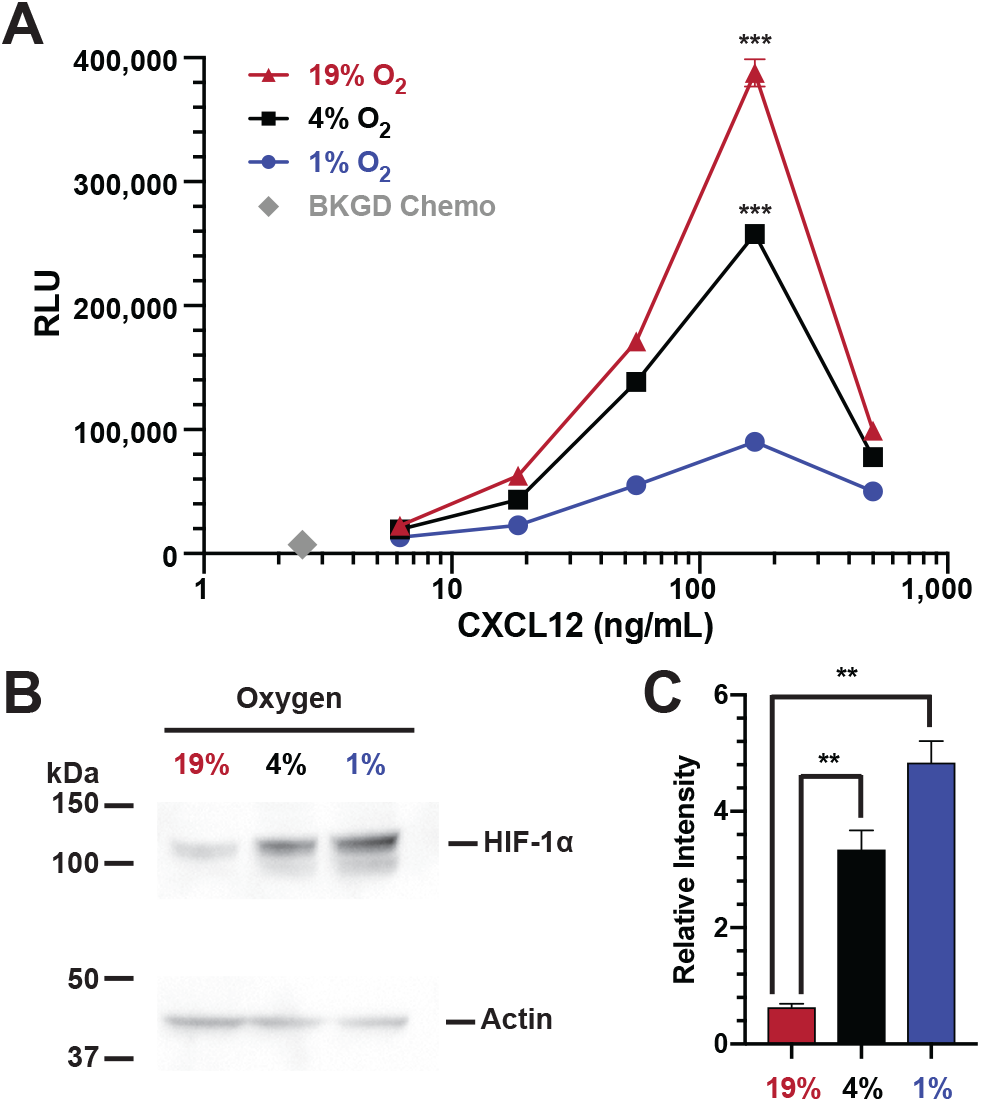
HIF-1*α* stabilization correlates with hypoxia induced CXCR4 hypo-responsiveness of the RAMOS B cell line. RAMOS B cells were incubated at the indicated O_2_ levels for 24 hours and then subjected to (**a**) chemotaxis assay or (**b**) lysates were prepared and westerns performed to measure HIF-1*α* levels with actin being used as a loading control. Shown is a representative blot. (**c**) Quantitation of relative HIF-1*α* levels from three separate experiments. \*\*\**P<*0.001; \*\**P<*0.01

We next confirmed this O_2_ effect in freshly isolated B cells from human peripheral blood and murine spleen. Primary human peripheral blood B cells (n=3 donors) incubated at 19% O_2_ for 24 hours migrate in responses to CXCL12 (Figure 2A). Importantly, decreased migration occurred at both 1% and 4% O_2_, similar to that observed in the RAMOS cell line, confirming that primary human B cell migration is inhibited at lower O_2_ tensions. Mouse B cells exhibited an identical effect of lower O_2_ levels on CXCL12 induced B cell chemotaxis (Figure 2B), with robust chemotaxis to CXCL12 at 19% O_2_, which decreases with 24 hours of incubation at 1% or 4% O_2_. Virtually identical CXCR4-CXCL12 chemotaxis in mouse and human B cells suggests this is a highly conserved regulatory system.

**Fig. 2.**
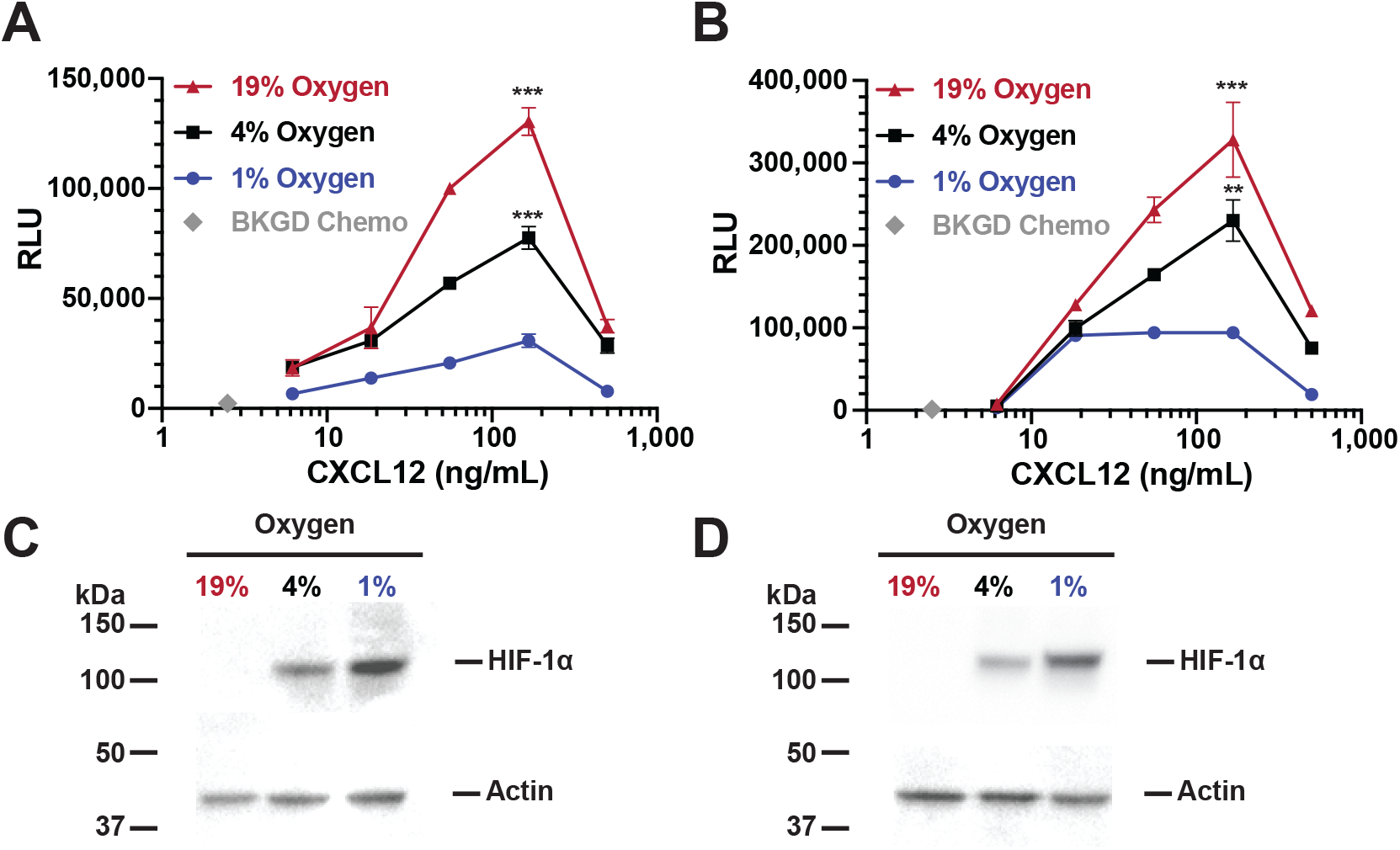
HIF-1*α* stabilization correlates with hypoxia induced CXCR4 hypo-responsiveness of primary human and mouse B cells. Isolated primary human peripheral blood B cells (**a**) or splenic mouse B cells (**b**) were incubated for 24 hours at the indicated O_2_ levels and then subjected to chemotaxis assay. Shown is representative experiment from 3 independent experiments for each B cell source. Lysates from either human (**c**) or mouse (**d**) B cells were prepared and westerns performed to measure HIF-1*α* levels with actin being used as a loading control. Shown is a representative blot.

Sensitivity of CXCR4 to CXCL12 is modulated by stabilization of the transcription factor HIF-1*α*, which occurs at low O_2_ levels in human B cells and induces nuclear translocation (5). To confirm HIF-1*α* stabilization, we performed western blot analysis of HIF-1*α* protein levels in nuclear lysates from human B cell lines and peripheral blood B cells, and from mouse splenic B cells. Incubation of the human RAMOS B cell line for 24 hours at 19% O_2_ results in barely detectable levels of HIF-1*α* protein (Figure 1B and C). In contrast, increasing levels of HIF-1*α* protein over time are detected as the cells are incubated at lower O_2_ levels (4% and 1%, a 5.3 and 7.6 fold increase as compared to 19%, respectively), demonstrating O_2_ dependent HIF-1*α* protein stabilization and nuclear localization. An identical finding was present in primary human B cells (Figure 2C). Similarly, mouse B cells incubated at 19% O_2_ do not exhibit detectable levels of HIF-1*α* protein in nuclear lysates (Figure 2D). Thus there is a negative correlation between nuclear HIF-1*α* protein levels and human or mouse B cell chemotactic activity.

In addition to HIF-1*α* stabilization, HIF-2*α* has also been shown to be stabilized upon hypoxic exposure of several different cell types (46, 47), but there is little data in human B cells. We performed a western blot analysis of HIF-2*α* protein stabilization in human B cells under varying O_2_ conditions. We did not observe any significant HIF-2*α* stabilization in human B cell lysates at low O_2_ levels (Supplementary Figure 2). These results suggest that HIF-2*α* does not play a significant role in O_2_ sensitive human B cell migration within the time frame our experiments are performed.

### HIF-1*α* is necessary to induce hypoxia dependent CXCR4 hypo-responsiveness

We hypothesized that HIF1*α* expression is necessary for B cell chemotactic hypo-responsiveness at low O_2_ levels, and evaluated this hypothesis using shRNA silencing of HIF-1*α*. Stable RAMOS B cell lentiviral transfectants were generated expressing either HIF-1*α* shRNA, or a control non-specific shRNA vector and evaluated at the peak chemotaxis dose of CXCL12 (166 *µ*g/mL; Figure 1A). RAMOS cells expressing HIF-1*α* shRNA had increased chemotactic activity at both 4% or 1% O_2_ levels as compared to either the control shRNA transfectants or non-transfected cells (Figure 3A). As predicted, HIF-1*α* protein levels were significantly decreased in HIF-1*α* shRNA transfected cells (Figure 3B and C). These genetic silencing results clearly demonstrate that HIF-1*α* expression is necessary for the observed decrease in CXCR4 responsiveness at low O_2_ levels.

**Fig. 3.**
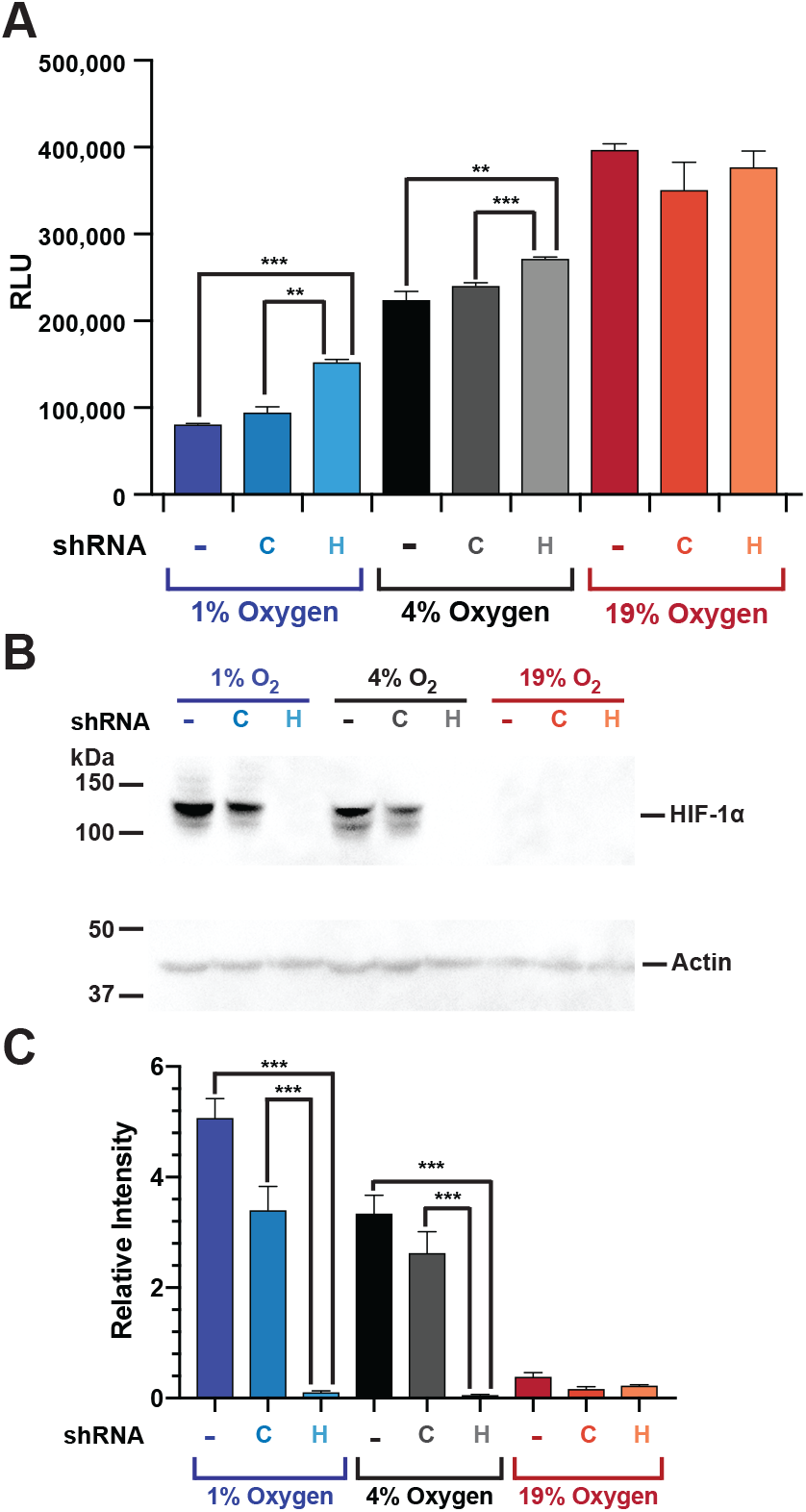
shRNA silencing of HIF-1*α* partially restores B cell CXCR4 responsiveness at low O_2_levels. RAMOS B cells, stably transfected with lentiviral vectors expressing either HIF-1*α* shRNA (H), a scrambled non-specific control sequence shRNA (C) or parental non-transfected RAMOS cells (-), were incubated at the indicated O_2_ levels for 24 hours and then subjected to (**a**) chemotaxis assay or (**b**) lysates were prepared and westerns performed to measure HIF-1*α* levels with actin being used as a loading control. Shown is a representative blot. (**c**) Quantitation of relative HIF-1*α* levels from three separate experiments. \*\*\**P<*0.001; \*\**P<*0.01

### Cyclosporine A inhibits hypoxia dependent CXCR4 hypo-responsiveness by destabilizing HIF-1*α*

Cyclosporine A (CyA) destabilizes HIF-1*α* in glioma cells (35) and we hypothesized that CyA could disrupt B cell HIF-1*α* stabilization, altering CXCL12 induced B cell migration. Consistent with this hypothesis, and the above data, exposure of the RAMOS B cell line to 1% O_2_ inhibits CXCL12 mediated B cell chemotaxis relative to that observed at 19% O_2_. However, treatment with increasing concentrations of CyA progressively increases CXCL12 mediated chemotaxis of RAMOS cells at 1% O_2_ (Figure 4A). These results are not restricted to cell lines as both primary human and mouse B cells also exhibit a similar increase in chemotactic activity after treatment with CyA (Figure 4D and E). In addition, the levels of CyA used here are physiologically relevant as serum levels of 10-2000 ng/mL are achieved *in vivo* in immunosuppressed renal transplant recipients (48).

**Fig. 4.**
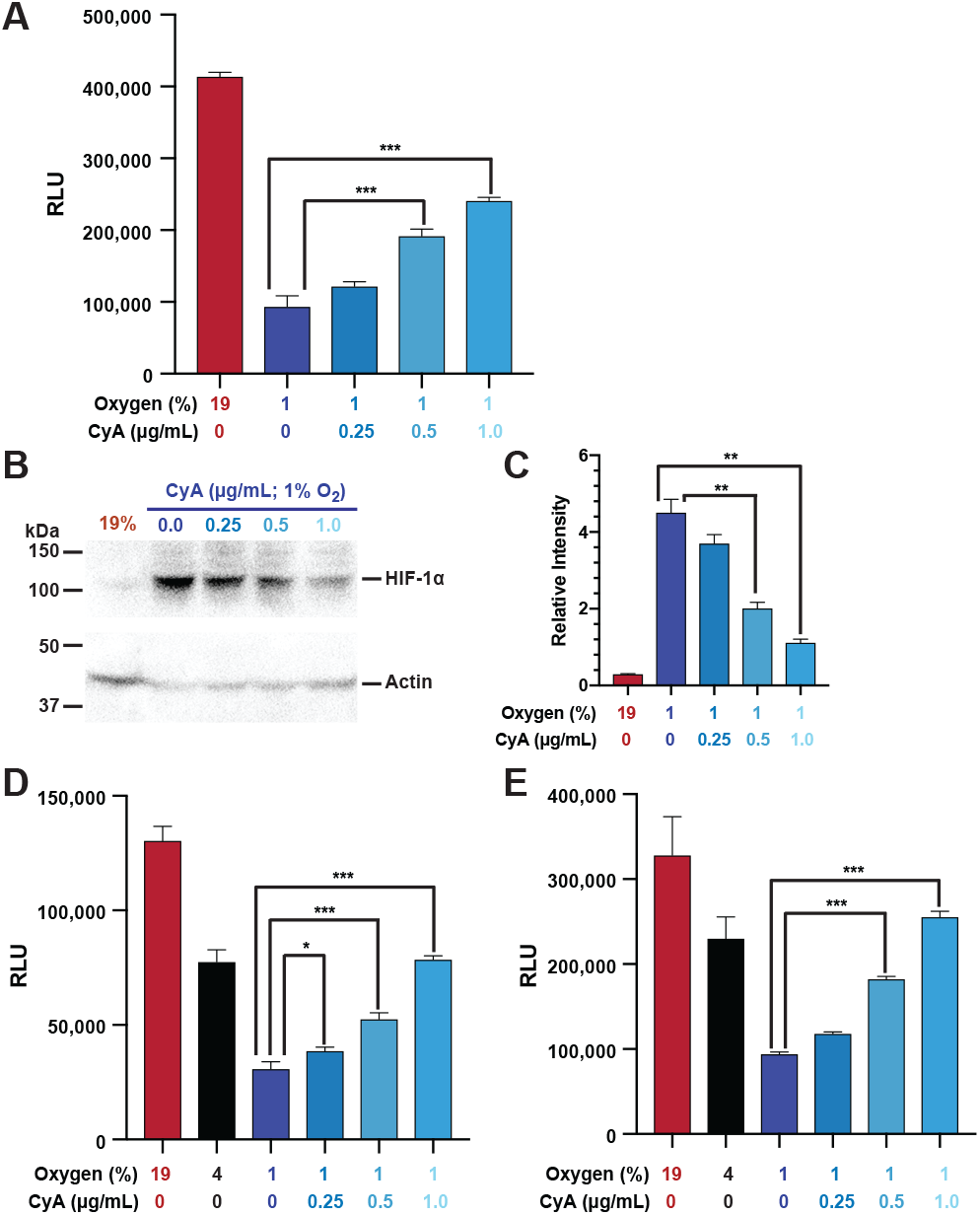
Cyclosporine A interferes with B cell hypoxia dependent CXCR4 hypo-responsiveness by destabilizing HIF-1*α*. RAMOS cells were incubated at 19, 4 or 1% O_2_ levels, with some 1% samples being incubated with the indicated concentrations CyA. After 24 hours the cells were subjected to (**a**) chemotaxis assay or (**b**) lysates were prepared and westerns performed to measure HIF-1*α* levels with actin being used as a loading control. Shown is a representative blot. (**c**) Quantitation of relative HIF-1*α* levels from three separate experiments. Primary human (**d**) or mouse (**e**) B cells were incubated at 19, 4 or 1% O_2_ levels, with some 1% samples being incubated with the indicated concentrations CyA. After 24 hours a chemotaxis assay was performed \*\*\**P*<0.001;\*\**P*<0.01;\**P*<0.05

To assess the correlation between CyA’s effect on migration and HIF-1*α* stabilization, we performed western blot analysis on RAMOS nuclear lysates. As is evident from Figure 4B and C, treatment of RAMOS cells, with increasing amounts of CyA at 1% O_2_ leads to a progressive decrease in HIF-1*α* levels. These results again directly correlate HIF1*α* with chemotactic activity and demonstrate that CyA treatment results in decreased HIF-1*α* protein levels, resulting in increased CXCR4 responsiveness, despite the fact that the B cells are incubated at low, 1% O_2_ levels.

### Pharmacological stabilization of HIF-1*α* at 19% O_2_ mimics the effect of incubating B cells at low O_2_

The above data defines an O_2_ sensing molecular switch controlling B cell migration that correlates with and requires HIF-1*α* stabilization, and is sensitive to CyA treatment. However, this does not rule out the possibility that other O_2_ sensitive factors independent of HIF-1*α* are responsible for the observed chemotactic effects. To address this, we performed experiments stabilizing HIF-1*α* at 19% O_2_, using the prolyl-4-hydroxylase (PHD) inhibitor dimethyloxallyl glycine (DMOG), thus eliminating potential side effects associated directly with low O_2_ levels. DMOG appears to selectively stabilize HIF-1*α*, with minimal effects on HIF-2*α* (47). We found that incubation of RAMOS cells with DMOG at 19% O_2_ results in decreased CXCL12 responsiveness (Figure 5A), similar to the effect observed incubating these same cells at low O_2_ levels. DMOG also has a similar effect on primary human and mouse B cells (Figure 5C and D). Western blot analysis (Figure 5B) of nuclear lysates from the RAMOS cell line confirms that the CXCL12 chemotactic effects highly correlates with HIF-1*α* protein stabilization, even when the cells were incubated at 19% O_2_.

**Fig. 5.**
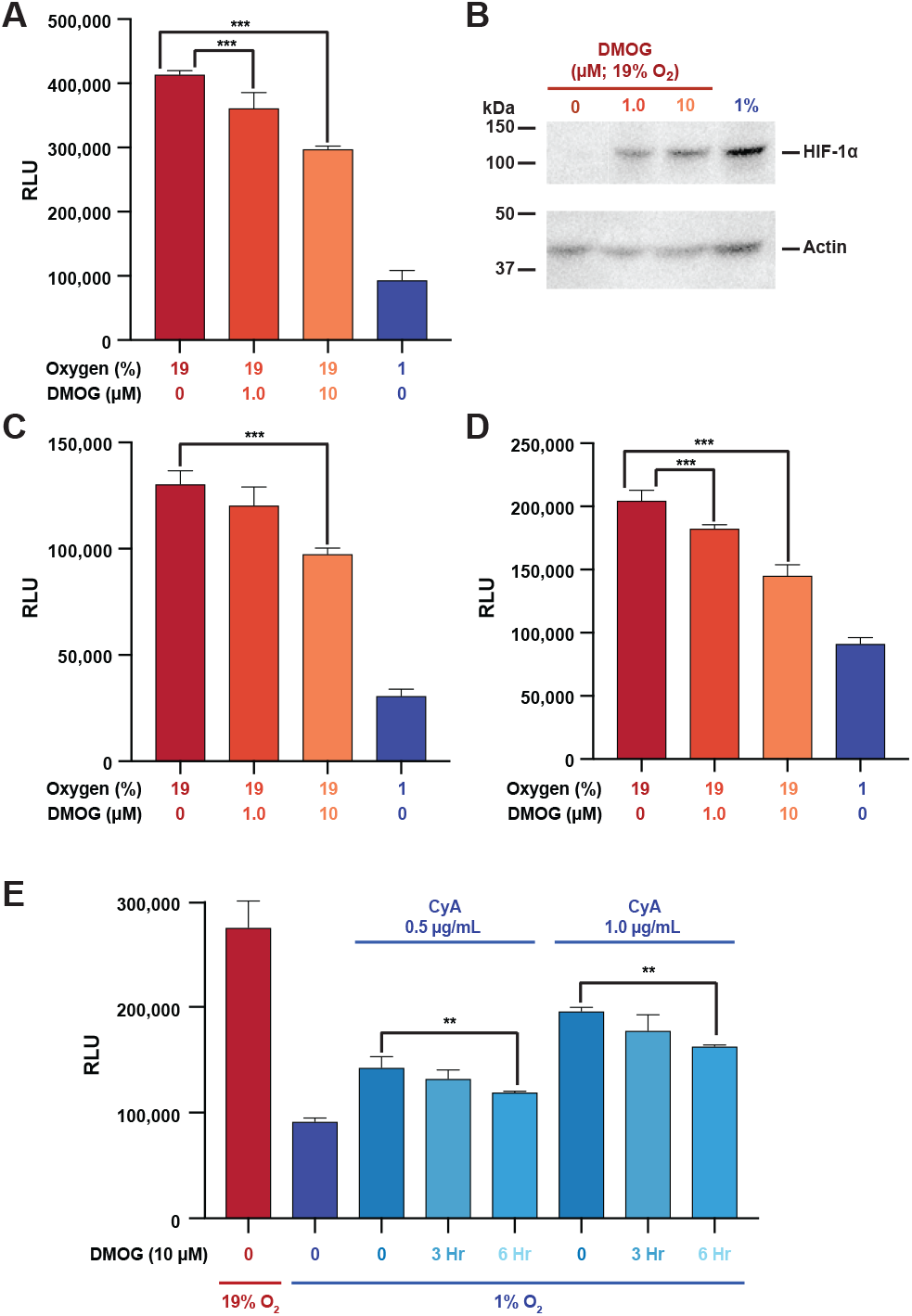
DMOG treatment of B cells stabilizes HIF-1*α* and partially restores CXCR4 hypo-responsiveness in the presence of Cyclosporine. **A.** RAMOS B cells (**a**), primary human peripheral blood B cells (**c**) or mouse plenic B cells (**d**) were incubated at 19% or 1% O_2_ levels for 24 hour with the indicated concentration of DMOG and then subjected to chemotaxis assay. (**b**) RAMOS cells lyates were assessed by western blot for HIF-1*α* levels with actin being used as a loading control. Shown is a representative blot. (**e**) RAMOS B cells were incubated for 24 hours at either 19% or 1% O_2_levels for 24 hours with the indicated concentration of CyA. Three or 6 hours before the end of the 24 hour incubation, DMOG was added to a final concentration of 10 *µ*M and then the cells were subjected to chemotaxis assay. \*\*\**P<*0.001; \*\**P<*0.01

### Transient restabilization of HIF-1*α* at 1% O_2_ during CyA treatment partially restores CXCR4 hypo-responsiveness

We have shown that CyA disrupts low O_2_ dependent CXCR4 hypo-responsiveness in B cells, in part through HIF-1*α* destabilization. We also show that pharmacological stabilization if HIF-1*α* at 19% O_2_ levels mimics the effect of low O_2_. We next asked whether transient DMOG addition at 1% O_2_ levels during CyA treatment might, at least partially, restore O_2_ dependent CXCR4 hypo-responsiveness. To test this hypothesis, we treated RAMOS B cells with CyA for 24 hours, and during the last 3 or 6 hours of the incubation, DMOG was added. Treated cells were then subjected to our chemotaxis assay. This transient DMOG treatment partially restored CXCR4 hypo-responsiveness, despite the presence of CyA (Figure 5E).

### Proteomics reveals HIF-1*α* regulated proteins and their restoration by CyA

To evaluate oxygen-dependent regulation of signaling components and their responsiveness to CyA, unbiased proteomics of RAMOS cell lysates at 19% and 1% O_2_ levels, and compared these results to those obtained at 1% O_2_ levels after CyA exposure. Figure 6A shows 15 proteins that responded to the change in oxygen levels, differentially expressed between cells at 19% oxygen and cells at 1% oxygen without CyA treatment. Of these, 6 had increased protein levels at 19% O_2_ and 9 were up-regulated under hypoxic conditions. These 9 up-regulated proteins included several regulated by HIF-1*α* such as BCL2 Interacting Protein 3 (BNIP3), EGLN1 (Egl nine homolog 1), Protein FAM162A (FAM162A), Prolyl 4-hydroxylase subunit alpha-1 (P4HA1), Fructose-bisphosphate aldolase C (ALDOC), 6-phosphofructo-2-kinase/fructose-2,6-bisphosphatase (PFKB1) and Procollagen-lysine,2-oxoglutarate 5-dioxygenase 1 (PLOD1). PLOD1 is a lysyl hydroxylase which regulates extracellular matrix stiffening and collagen fiber alignment along with prolyl 4-hydroxylase subunit alpha-1 P4HA1 and HIF-1*α* (49). FAM162A (50) and BNIP3 (51) promote intrinsic apoptosis and autophagy in response to hypoxia via interactions with HIF-1*α*. Moreover, HIF-1*α* coordinately represses and activates genes such as ALDOC and PFKB1 to effect metabolic switching to a glycolytic state, providing biosynthetic substrates that facilitate proliferation (52).

**Fig. 6.**
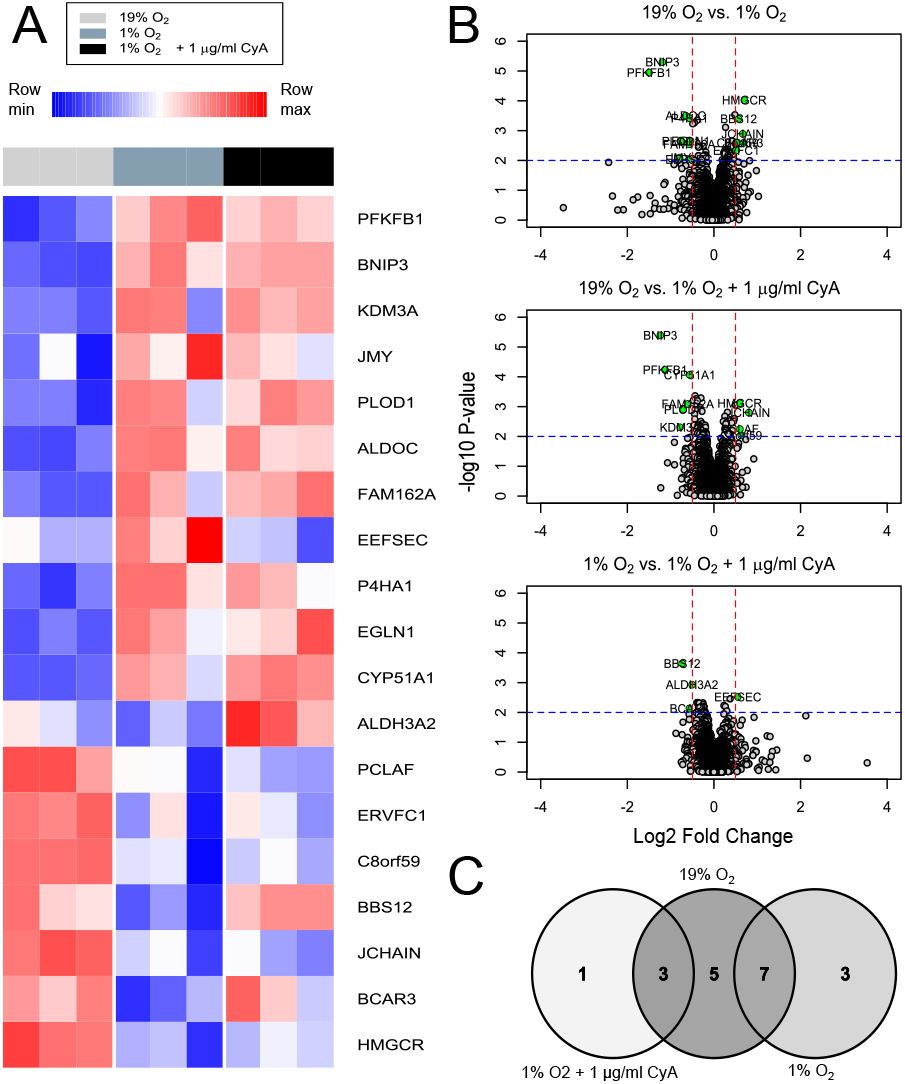
Differential analysis of proteomics data. (**a**). Heatmap of proteins with p-value ≤ 0.01 and log2FC ≥ 0.5 in any contrast. Color in each tile represents the scaled abundance value. (**b**). Volcano plots of proteins in the contrasts 19% O_2_ vs. 1% O_2_, 19% O_2_ vs 1% O_2_ + 1 *µ*g/mL CyA, and 1% O_2_ vs 1% O_2_ + 1 *µ*g/mL CyA. (**c**). Venn diagram of proteins with differential abundances from the three contrasts.

We were particularly interested in proteins that were restored after treatment with CyA at 1% oxygen 6C. 7 out of 18 proteins did not respond differently to the hypoxic conditions when the cells were treated with CyA. The treatment with CyA restored the abundance of 8 proteins at 1% oxygen to a level similar to that in cells grown at 19% oxygen. Closer examination of these proteins reveals three candidates, selenocysteine-specific elongation factor (EEFSEC), Bardet-Biedl syndrome 12 protein (BBS12) and breast cancer anti-estrogen resistance protein 3 (BCAR3), that could be involved in the regulation of CXCR4 responsiveness. EEFSEC, BBS12 and BCAR3 have been shown to be responsive to cellular oxygen metabolism (53–55). Pathway analysis of all differentially expressed proteins revealed the role of 9 pathways linked to regulation of HIF-1*α* and glucose metabolism in restoring the responsiveness to oxygen levels upon CyA treatment. Thus differential analysis revealed several signaling components that could be dysregulated in RAMOS cells due to hypoxia and further controlled by CyA.

### Protein association networks identify signaling components perturbed by CyA

Differential analysis identified only 0.4% proteins to be responsive to the oxygen levels and/or CyA. To expand the search for relevant proteins we sought to investigate functionally related proteins both from (a) our dataset and (b) in the literature. We constructed a co-expression network in which high degree nodes are involved in several metabolic processes Figure 7A. For example N6-adenosine-methyltransferase non-catalytic subunit (METTL14) is required in hypoxic stabilization of mRNAs (56) This is consistent with the known role of HIF-1*α* in regulating metabolic switches.

**Fig. 7.**
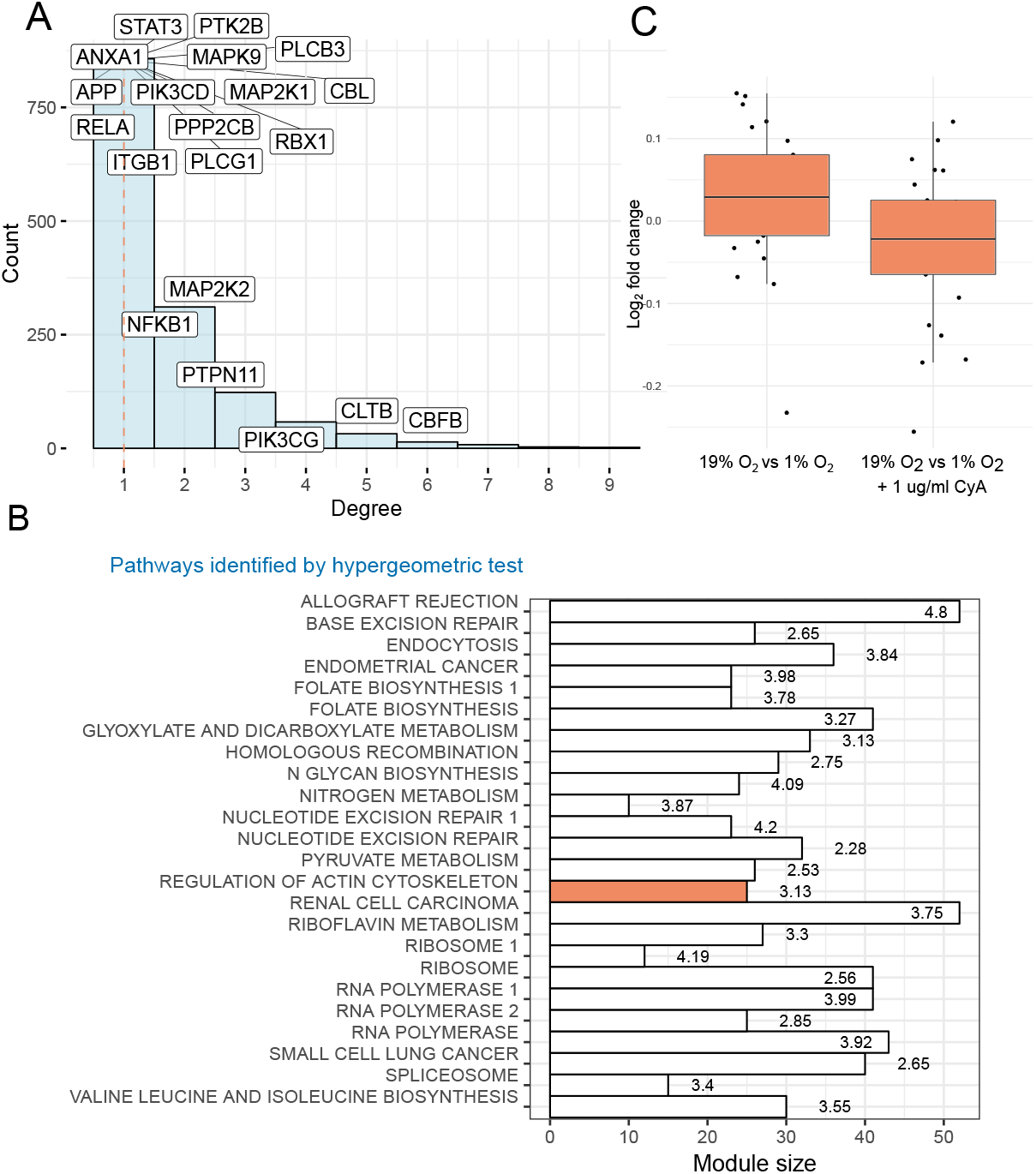
Protein association networks identify CXCR4-associated signaling components. (**a**). Distribution of node degrees in the co-expression network. 20 proteins with highest degree in knowledge based networks are highlighted. The dashed red line indicates median node degree. (**b**). Module sizes (of size ≥ 10) identified from the co-expression network (x-axis) across the top enriched pathway identified by the hypergeometric test (p ≤ 0.05) are shown. Text labels show -log_1_ 0 p-value from the hypergeometric test. The red bar indicates the only module with significantly different log_2_ fold change in 19% *O*_2_ vs 1% *O*_2_ and 19% *O*_2_ vs 1% *O*_2_ + CyA contrasts (*p* ≤ 0.05, two-tailed t-test) (**c**). Distribution of *log*_2_ fold changes of the proteins (medians across three replicates) in the labeled module in 19% *O*_2_ vs 1% *O*_2_ and 19% *O*_2_ vs 1% *O*_2_ + CyA contrasts.

We next compared the protein co-expression network with a knowledge-based network constructed using the STRING database (43). We found overlap between 72 nodes present in the knowledge-based and co-expression networks. Figure 7A shows the top 20 proteins, sorted by node degree, and overlaid on a histogram of the degree distribution of the co-expression network. Several high-degree nodes from the knowledge-based network (e.g. core-binding factor subunit beta (CBFB), clathrin light chain B (CLTB), phosphatidylinositol 4,5-bisphosphate 3-kinase catalytic subunit gamma isoform (PIK3CG)) were also highly-connected in our co-expression network, suggesting their key role in the regulation of HIF-1*α* and CyA signaling. Core-binding factor subunit beta is a master-regulator of genes involved in hematopoiesis (e.g., RUNX1) and osteogenesis (e.g., RUNX2). Clathrin light chain B is involved in receptor-mediated endocytosis (57).

To identify modules of highly associated proteins, and by inference their genes, we used the edge-betweenness commnity detection algorithm (40). We identified 24 communities with ≥10 proteins. These modules were characterized by the significantly enriched pathways (Figure 7B). One community was of particular interest since it was restored after treatment with CyA at 1% oxygen Figure 7C and was enriched for proteins involved in regulation of the actin cytoskeleton (see Supplementary Table 1 for a list of the top 3 enriched pathways, with associated p-values from the hy-pergeometric test). Specifically, this module contains the proteins PFN2 (profilin 2), IQGAP2 (Ras GTPase-like activating protein), and ITGAL (integrin alpha L). Both PFN2 (58) and IQGAP2 (59) regulate cell motility in response to intracellular signals. PFN2 is an epigenetic regulator of SMAD2/3 (Mothers against decapentaplegic homolog 2/3) (60), which in turn has been shown to promote metastasis in cancer cells (61). SMAD2 and SMAD3 are also regulated by HIF-1*α* (62). In the knowledge-based STRING network, SMAD3 is directly linked to HIF-1*α*. ITGAL is a target of the immunosuppressant drug efalizumab, which acts by reducing lymphocyte migration to sites of inflammation (63). Thus, these proteins may be involved in the restoration of B cell chemotactic activity under hypoxic conditions by CyA as measured by our chemotaxis assay and suggest targets for further investigation.

## Discussion

The results we report here, characterization of an O_2_ sensing-HIF-1*α* dependent molecular switch that alters human and mouse B cell CXCR4 responsiveness, which is disrupted by the calcineurin inhibitor cyclosporine A (CyA), have broad implications in regards to normal immunological processes. These processes occur naturally at differing O_2_ levels *in vivo*, an aspect of immunological function that is often overlooked, and the particular importance of O_2_ levels in B cell specific responses has only recently begun to come to light. For instance, it has been shown that GC hypoxia itself is important for efficient B cell class switching, plasma cell differentiation and abundant antibody production (16). In addition, it has also recently been reported that the hypoxia within the GC light zone is critical for generation of high quality antibody responses in part by creating a more stringent threshold for vital survival signals, leading to the selection and survival on higher affinity antibody producing B cells (64). Also, CXCR4 signaling has been shown to be critical for normal B cell function and development (29, 31), and specifically, proper GC B cell localization within the light or dark zone is necessary (65), and this localization has been shown to be dependent on CXCR4 signaling (27). Importantly, GC B cells, which reside within a highly hypoxic environment, have actually been shown to express surface CXCR4, but they are unresponsive to CXCL12 (33). Increasingly hypoxia has been connected to CXCR4 regulation and function (32, 33).

Combined, these results link proper B cell GC localization, through differential CXCR4 responsiveness controlled by O_2_ levels, to optimum B cell responses. In direct support, we define here a HIF-1*α* dependent O_2_ sensing molecular switch that controls the responsiveness of CXCR4 to CXCL12, independent of CXCR4 surface expression, directly linking GC hypoxic conditions to CXCR4 signaling. Importantly, we show that CXCR4 responsiveness is modulated through O_2_ levels that are physiological observed within the LN. In particular, LN O_2_ levels vary from 4-5% O_2_ near afferent vessels to <1% in portions of the GC (16, 17, 33, 66). Thus, it is very likely that this process is naturally coordinated and necessary for appropriate nodal and GC B cell dark/light zone localization and any disruption to this process would likely result in substandard B cell responses.

Calcineurin inhibitors (CNI) are commonly utilized to ameliorate immunological allograft rejection in solid organ transplant recipients.[37-39] However, the overall immunosuppressive effects of CNI results in decreased antibody production, which can lead to greater morbidity and mortality associated with common infectious agents, such as influenza viruses (67–69). One means of protecting these vulnerable populations is through vaccination, however, the immunosuppressive nature of CNI also leads to suppressed vaccine responses (2, 3). Currently, CNI dependent inhibition of NFAT in helper T cells, leading to decreased T cell proliferation and IL-2 production, is largely believed to account for these decreased antibody responses (4). Despite these T cell effects, direct CNI effects on B cells have remained largely unidentified. We show here that CyA has a novel and direct effect on B cell migration by disrupting the natural O_2_ sensing molecular switch we have defined by destabilizing HIF-1*α*, allowing B cells to preserve CXCR4 responsiveness at low (<1%) O_2_ levels. Preservation of CXCR4 responsiveness would likely disrupt coordinated GC B cell dark zone vs. light zone localization, potentially leading to suppressed B cell responses *in vivo*. This is the first time, to our knowledge, that CyA have been shown to directly affect B cell migration, identifying a novel, and targetable pathway by which CyA may directly affect B cell responses.

Unbiased proteomics analysis identified signaling components involved in CXCR4 and HIF-1*α* signaling that are modulated by CyA. The differential expression analysis identified signaling pathways responding to hypoxia and their key regulators. Several of the identified pathways are associated with HIF-1*α*, in consensus with the previous literature (70–73). However, several of the proteins reversing the effect of hypoxia with drug had lower fold-changes upon treatment with CyA. Hence, we expanded our analysis by developing association networks. The network analysis allowed us to probe CXCR4 and HIF-1*α* signaling specifically even when CXCR4 expression itself was not changed. The network analysis revealed key regulators involved in the response to CyA and hypoxia. Moreover, it identified a module which reversed the effect of hypoxia upon CyA treatment and revealed a novel, highly intra-correlated protein module potentially involved in the responsiveness to CyA.

Improvement of immune function in CyA treated transplant patients during vaccination has the potential to increase vaccine efficacy, resulting in decreased morbidity and mortality associated with infection. However, as immune suppression is required to avoid allograft rejection, such immunological improvement would have to be transient and applied only during vaccine immune responses. We have identified a potential pathway to target, namely, reversal of CyA effects by transiently re-stabilizing HIF-1*α*. Indeed our *in vitro* data demonstrates that transient re-stabilization of HIF-1*α* through treatment of CyA affected B cells with the prolyl-4-hydroxylase (PHD) inhibitor dimethyloxallyl glycine (DMOG) results in retention of O_2_ dependent changes in CXCR4 responsiveness. It is important to note that PHD inhibitors are clinically available for study, including FG-4592 and GSK1278863. However, preclinical study in mouse models would be required, and our data clearly demonstrates that both changes on O_2_ levels, as well as the CyA identically affect the migratory capacity of mouse B cells as compared to human B cells, allowing for direct *in vivo* preclinical studies.

## Conclusions

We have characterized an O_2_ sensing-HIF-1*α* dependent molecular switch that alters human and mouse B cell CXCR4 responsiveness. This switch likely plays a significant role in GC B cell development and function. We have also identified a novel and direct CyA B cell affect whereby CyA directly interferes with this molecular switch by destabilizing HIF-1*α* at low (<1%) O_2_ levels, preserving both human and mouse B cell CXCR4 responsiveness. The clinical implications of these results are potentially profound, as they identify a readily targetable pathway, through the transient use of clinically available PHD inhibitors (e.g. FG-4592 and GSK1278863), which may improve vaccine responses in vulnerable immune suppressed transplant patients.

## ABBREVIATIONS

AF: ammonium formate
AGC: automatic gain control
BCA: bicinchoninic acid
CD: cluster of differentiation
CNI: Calcineurin inhibitors
CyA: cyclosporine A
DMOG: dimethyloxallyl glycine
DTT: dithiothreitol
ECL: enhanced chemiluminescence
EDTA: ethylenediaminetetraacetic acid
FAK: Focal adhesion kinase
FBS: fetal bovine serum
FDR: false discovery rate
GC: Germinal center
HCD: higherenergy collisional dissociation
HIF: Hypoxia inducible factor
HPLC: high performance liquid chromatography
IL-2: Interleukin 2
HRP: horseradish peroxidase
LN: Lymph node
IMDM: Iscove’s modified Dulbecco’s medium
MAPK: Mitogenactivated protein kinase
MKP-1: MAP kinase phosphatase 1
NFAT: Nuclear factor of activated T-cells
O/N: Overnight
PBMC: peripheral blood mononuclear cell
PBS: phosphate-buffered saline
PHD: prolyl-4-hydroxylase
PVDF: polyvinylidene difluoride
RBCs: Red blood cells
RGS1: Regulator Of G Protein Signaling 1
RLU: relative luminescent units
RPMI: Roswell Park Memorial Institute
RT: room temperature
SCS: single cell suspension
shRNA: Small hairpin ribonucleic acid
TBS: tris-buffered saline
TBS-T: TBS with tween-20
TEAB: triethylammonium bicarbonate
TFA: trifluoroacetic acid
TMT: tandem mass tag

## ACKNOWLEDGEMENTS

The authors would like to thank Alexander Wiltse and Jason Emo for their excellent technical assistance, and Jiong Wang for her critical reading of the manuscript.

## DATA AVAILABILITY

Data for all experiments are available from the authors upon request. Please contact the corresponding author for more information. The remainder of the reagents are available publicly, as referenced in the Methods section.

## AUTHOR CONTRIBUTIONS

SPH conceived of the research, performed the research, and wrote the manuscript. MGP performed a portion of the analysis. DL performed a portion of the analysis. JG performed the research. VC performed the research. JT conceived of the proteomics analysis, and wrote the manuscript. MSZ conceived of the research and wrote the manuscript.

## ETHICS APPROVAL AND CONSENT TO PARTICIPATE

This study was approved by the Research Subjects Review Board at the University of Rochester Medical Center. (RSRB Protocol 71460), and all subjects gave written informed consent as described in the Methods section above.

## Supplementary Material

**Supplementary Figure 1.**
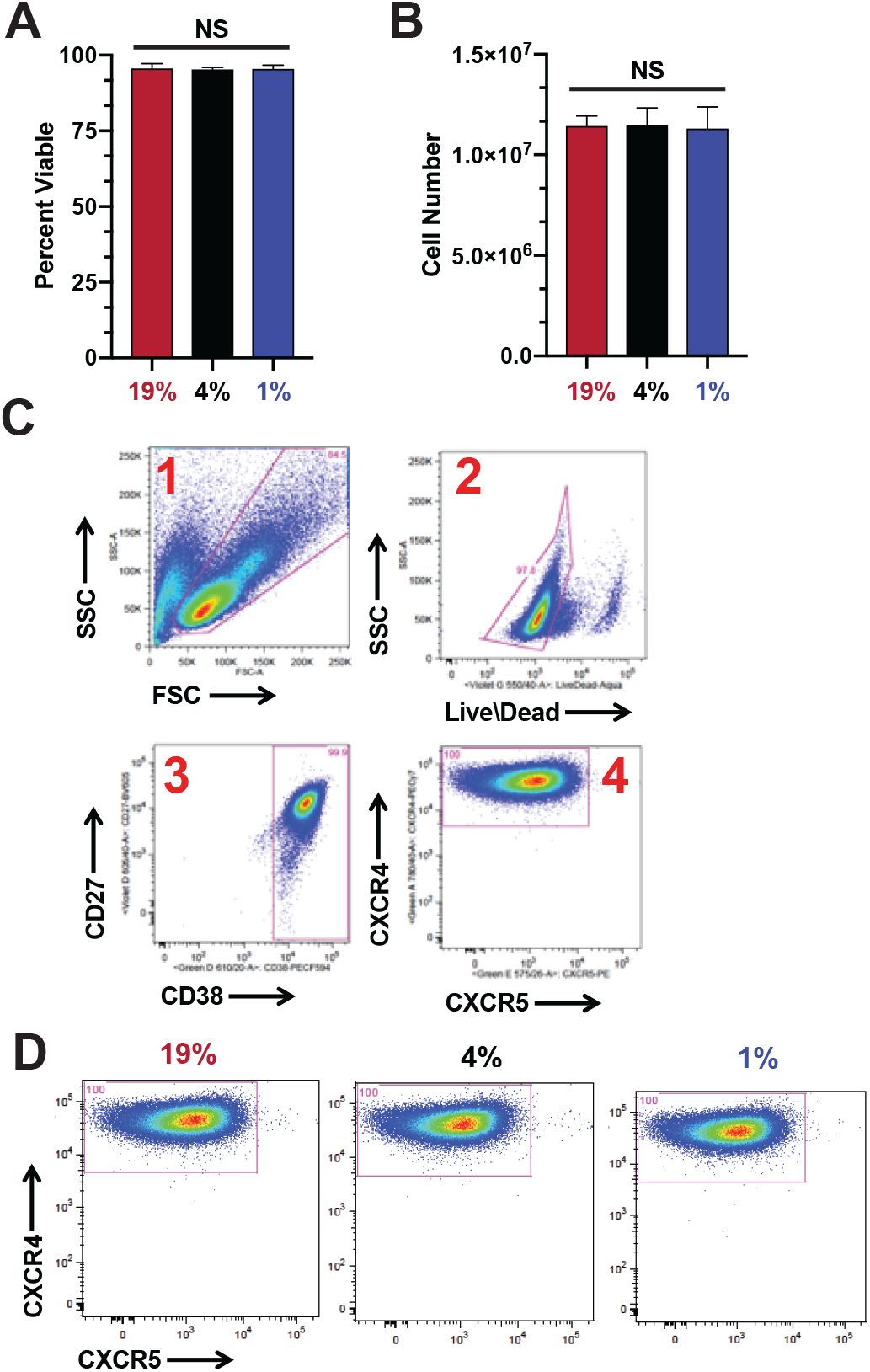
Chemotactic differences are not due to changes in cellular viability, proliferation or CXCR4 surface expression. RAMOS cells (5 × 10^6^ cells per condition at 5 × 10^5^ cells/mL, in triplicate) were incubated at the indicated O_2_ levels for 24 hours. After the incubation, cells were harvested and assessed for (**a**) viability (trypan blue exclusion), (**b**) proliferation by assessing total cell numbers or (**c-d**) surface CXCR4 levels assessed by flow cytometry. (**c**) Sequential gating strategy. (**d**) CXCR4 surface staining does not significantly change with changing O_2_ levels. Shown is representative experiment from 3 independent experiments. NS = not significant.

**Supplementary Figure 2.**
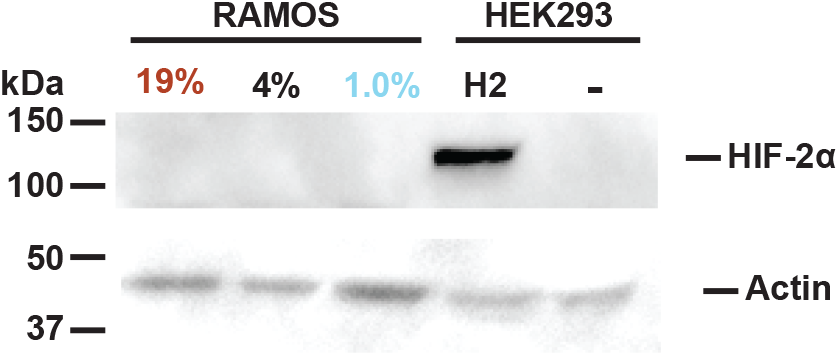
HIF-2*α* stabilization is not observed in RAMOS B cells after exposure to low O_2_ levels. RAMOS cells were incubated at the indicated O_2_ levels for 24 hours. After the incubation, cells were harvested, lysates were prepared and westerns performed to measure HIF-2*α* levels with actin being used as a loading control. Lysates from HIF-2*α* over-expressing (H2) or non-transfected (-) HEK293 cells, were used as positive and negative controls, respectively. Shown is a representative blot from 3 independent experiments.

**Supplementary Figure 3.**
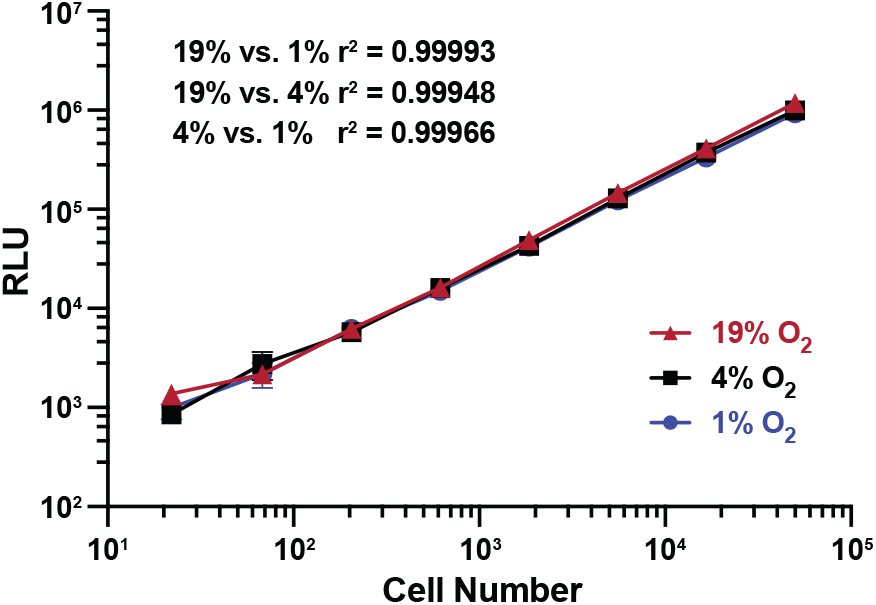
A direct correlation exits between cell numbers and RLU readout regardless of O_2_ levels. RAMOS cells (5 × 10^6^ cells per condition at 5 × 10^5^ cells/mL, in triplicate) were incubated at the indicated O_2_ levels for 24 hours. After the incubation, cells were harvested and counted. Cells were then plated in triplicate well in 96 well plates at the indicated cell numbers. Plates were then incubated at the original O_2_ levels for an additional hour and luminescence was then assessed to mimic the conditions of the chemotaxis assay. assessed for (**a**) viability (trypan blue exclusion), (**b**) proliferation by assessing total cell numbers or (**c**) surface CXCR4 levels assessed by flow cytometry. Shown is representative experiment from 3 independent experiments. Pearson correlation coefficients were calculated using Prism software (GraphPad Software, San Diego, CA; Mac version 8.0.2)

